# Lifestyle modulates riverbed diazotrophy: from abundance and activity to diversity of free-living and biofilm-associated N_2_ fixers

**DOI:** 10.64898/2026.04.27.721245

**Authors:** Rand Lander, Carla Villamarin, Hagar Siebner, Shai Arnon, Edo Bar-Zeev

**Affiliations:** Zuckerberg Institute for Water Research, The Jacob Blaustein Institutes for Desert Research, Ben-Gurion University of the Negev, Sede Boqer Campus, 8499000, Israel

**Author notes:** Corresponding author: Edo Bar-Zeev,;.

**Keywords:** N_2_ Fixation, Diazotroph, Freshwater, Streambed, Sediment, Biofilm, Hyporheic zone

## Abstract

Despite the critical role that biological N_2_ fixation plays in controlling primary and secondary production in aquatic ecosystems, freshwater subsurface diazotrophy has received minimal attention. Here, we quantified N_2_ fixation rates, diazotroph abundance, and diversity in the hyporheic zone across seasons and redox regimes, distinguishing between free-living and biofilm-associated lifestyles. Samples were collected from aerobic and suboxic strata of the Jordan River streambed and incubated in microcosms for 24 h with dissolved ^15^N_2_ under ambient conditions. Immunolabeling the nitrogenase enzyme coupled with flow-cytometry revealed that diazotrophs accounted for ≤9.6 % of total bacterial abundance but were consistently enriched in biofilms. These biofilms supported 377-fold higher N_2_ fixation rates per-cell than free-living diazotrophs. Confocal laser scanning microscopy captured thicker (several tens of microns) complexes of extracellular polymeric substances (EPS) encasing aerobic biofilms than suboxic, especially during winter. Biofilm-associated communities retained high abundances (≤21 %) of facultative and obligate anaerobes in oxic zones. Compared to those in biofilms, the abundance, N_2_ fixation rates, and diversity of the free-living fraction varied much more across both aerobic and suboxic zones. *nifH* sequencing unveiled the dominance of Pseudomonadota and Thermodesulfobacteriota across all communities. Overall N_2_ fixing activity, when converted to volumetric units, was 1.5 × 10^5^-fold greater in the streambed than the overlying water. These findings identify the hyporheic zone as an active hotspot for benthic diazotrophy and intimate the centrality of microbial lifestyle in determining how diazotrophs persist and remain highly active in fluctuating redox and nutrient conditions in the freshwater ecospace.

## Introduction

Diazotrophs are bacteria and archaea that possess a functional nitrogenase enzyme complex, encoded by the *nifH* gene, which facilitates the bioconversion of dinitrogen (N_2_) into biologically reactive molecules (^1^). N_2_ fixation is paramount to fueling the nitrogen budget of aquatic ecosystems, where varying fixation rates can impact primary and secondary production (^1^). Diazotrophs require an extensive supply of energy-rich molecules (∼16 ATP per N_2_ molecule reduced) to fix N_2_ into ammonia (^2,3^). Autotrophic diazotrophs can supply the organic requirements via photosynthesis (Luo et al., 2014) or chemosynthesis (^4^), while heterotrophic diazotrophs rely on external organic carbon sources (^5,6^). Regardless of their metabolic traits, diazotrophs are often impaired by dissolved oxygen, DO (^7^), high concentrations (≥1 µM) of dissolved inorganic nitrogen, DIN (^8^), low concentrations of phosphorus ([PO_4_] <5-42 μM) (^9^), and insufficient supply of key trace elements (e.g., Mo and Fe) (^1,10,11^). Diazotrophs can cope with adverse environmental conditions by modulating their lifestyle, alternating between free-living, tight associations with biofilms and aggregates (^12–16^), and symbiotic relationships with other organisms (^11^). Diazotrophy was predominantly studied in marine environments, with a severe paucity of research in the freshwater biosphere. Compared to marine ecosystems, the fluvial biospheres are highly heterogeneous with N_2_ fixation rates of 400 ± 70 nmol N L^-1^ d^-1^ in the water column and 2.4 ± 4 nmol N g^-1^ d^-1^ in the benthos (considering ^15^N_2_ enrichment assays only) (^17^). Yet, these studies focused on the water column and benthic environments, with little consideration of subsurface diazotrophy.

Subsurface sediments of fluvial ecosystems are often highly reactive near the water-sediment interface, where exchange of water occurs between the stream and sediments omnidirectionally (^18,19^). This area within streambeds may extend up to several tens of cm, depending on physicochemical conditions, and is herein defined as the hyporheic zone (^20^). The hyporheic zone is notably characterized by strong gradients of nutrients, organic matter, and dissolved oxygen (DO), resulting in swift transitions from aerobic to suboxic conditions (^20,21^). Throughout this subsurface zone, bacteria can be found as biofilms attached to the sediment, or as free-living cells in the pore water. Despite the potential importance of diazotrophs to nitrogen cycling, the abundance, N_2_ fixation rates, and diversity of diazotrophs in the subsurface of freshwater ecosystems remain largely unexamined.

In this study, subsurface diazotrophy was quantified through measurements of abundance, N_2_ fixation rates, and *nifH* community composition and diversity using core-microcosms collected from the Jordan River, Israel. Diazotrophs were quantified by flow cytometry following immunolabeling of the nitrogenase enzyme and spatially localized using confocal laser scanning microscopy. N_2_ fixation rates were determined with a ^15^N_2_ enrichment assay, and diazotrophs were identified by sequencing the *nifH* gene. Ultimately, this research provides comprehensive new insights into subsurface diazotrophy under the strong physicochemical gradients characteristic of this freshwater ecosystem, with a specific focus on free-living and biofilm-associated diazotrophic lifestyles.

## Materials and Methods

### Sampling Setup

Sediment cores and porewater were collected from the Jordan River at a downstream site (2—4 meters from the bank) during the summer and winter of 2024-2025 at depths of 1—20 cm (Figure S1). Porewater was extracted from the hyporheic zone using a custom-made, stainless-steel piezometer with thin slits (0.1 cm wide and 1 cm long) covered by a 55 μm mesh to filter out large particles. Oxic conditions in the depth profile of the subsurface were measured in real time by drawing porewater with a 60 mL syringe attached to the piezometer from different depths through a custom-made flow-through chamber equipped with a DO sensor (Xylem Analytics) (Figure S1B). The aerobic zone was defined at an oxygen saturation >85 %, while the suboxic zone was <40 % (Table S1). In addition, pH, electrical conductivity (EC), and temperature were measured on-site from the water column and porewater (Table S2). Additional porewater samples (500 mL) were collected from the aerobic and suboxic zones for measuring total organic carbon (TOC) and total nitrogen (TN). Sediment cores were then collected (100 mL of media) using clear PVC tubes driven into the aerobic and suboxic zones. The media from each core was added into 300 mL sterile, gas-tight, glass bottles with porewater from the same depth and kept in the dark. Six core-microcosms were collected from each depth within the aerobic and suboxic zones during each sampling campaign.

Altogether, there were two summer campaigns with n = 26 microcosms and three winter campaigns with n = 37 microcosms.

### Setup and sampling of the core-microcosms

Aerobic core-microcosms were uncapped for 12 h and suboxic samples were purged with N_2_ for 10 min to ensure DO saturation was below 15 %, then re-sealed. Three replicate microcosms were taken for both aerobic and suboxic zones to be enriched with 18 ml of ^15^N_2_ water (61 µM N final concentration), shaken to completely mix the porewater with the sediment, and incubated for 24 h under ambient conditions. Two replicate microcosms were taken from each aerobic and suboxic zone as natural abundance (not enriched with ^15^N_2_). The oxic conditions were further monitored by measuring DO with a probe in dedicated bottles incubated under the same conditions.

#### Sediment processing and biofilm disruption

At the terminus of the incubation period, the porewater and sediment were separated. The porewater from each core-microcosm was decanted into new, sterile 300 mL glass bottles. The residual sediment in each core-microcosm was suspended in 100 mL of saline water (0.4 % NaCl) containing 5 mM ethylenediaminetetraacetic acid (EDTA, Sigma Aldrich, Cat. No. 03690), which acts to dismantle the biofilm matrix. The suspensions were shaken vigorously and probe-sonicated (Qsonica Q55, 60 amp) three times for 30 seconds per cycle. The water-sediment slurry was allowed to settle for five minutes before collection or filtration.

### Analytical methods

#### ^15^N_2_ Stable Isotope Analysis

The supernatant containing cells detached from the sediment, were filtered through 25 mm pre-combusted (500 ^°^C, 4.5 h) glass fiber filters, GF/F, using a peristaltic pump. Volumes, ranging from 10—100 mL, were filtered to ensure sufficient nitrogen biomass for the δ^15^N measurements (>10 µg N and/or signal amplitude >1000 mV). Filters were then dried overnight at 60 °C and stored in a desiccator until further analysis. Dry samples were analyzed according to Geslier et al. (^6^). Calculations for heterotrophic N_2_ fixation were performed according to Mullholand et al. (^22^) and Geisler et al. (^6^). Additional details on sample preparation and the use of standards are provided in the supplementary information.

#### Cellular quantification of diazotroph and bacteria via flow cytometry

Porewater and sediment slurry were collected in triplicates (1.7 mL) from both aerobic and suboxic zones and fixed on-site with 6 μl of glutaraldehyde. Samples were then flash frozen in liquid nitrogen and stored at −80 °C until analysis. Before analysis, samples were thawed, incubated with 5 mM EDTA for 10 min and probe sonicated (30 s × 3 times). Diazotrophs were quantified after immunolabelling the nitrogenase enzyme in accordance with the method described by Geisler et al. (^14^), with minor modifications which are provided in the supporting information.

#### Visualization and microlocalization of diazotrophs in biofilms attached to sediment

Sediment particles (n = 10—20) were randomly collected from each aerobic and suboxic zone and placed onto a 1 μm polycarbonate filter (GVS, Life Sciences, USA) and gently filtered under vacuum (∼ 50 mbar). Particles were washed three times with 5 mL PBST. A primary antibody solution was added with 4.7 μl mL^-1^ anti-*nifH* and filtered PBST+BSA, then incubated for 1 h in the dark. After, it was washed again with PBST and incubated with a secondary antibody solution with 6 μl mL^-1^ Alexa Fluor^TM^ 488 goat anti-chicken and washed three times with PBS. Samples were stained with a 4 % Alcian blue solution (^23^) and 4’,6-diamidino-2-phenylindole solution (DAPI, 250 μg mL^-1^, Thermo Fisher Scientific D1306) for 45 min in the dark (^24^). Alcian blue and DAPI stains EPS and DNA, respectively. Excess stain was washed with 5 mL of PBS. Sediment particles were visualized within three days by a confocal laser scanning microscope (CLSM) from summer aerobic (n = 6) and suboxic (n = 7) biofilms, and winter aerobic (n = 14) and suboxic (n = 14) biofilms.

#### DNA extraction and nifH amplification

Prior to DNA extraction, porewater was filtered *in-situ*, through a Sterivex (C3235) filter until clogged. Sediment samples were taken directly from cores. The Sterivex and sediment samples were kept on ice and in the dark after sampling, then frozen in −80 °C until analysis within a few days. For sediment, the EDTA and NaCl slurry was made as described above, centrifuged at 1000 rpm for five minutes, and finally filtered through the Sterivex filter. DNA was extracted from the filters using the PowerWater Sterivex Kit (QIAGEN). Amplicon sequencing was performed in an Illumina MiSeq platform at the Genomics and Microbiome Core Facility (Rush University, Chicago, USA). Library preparation was accomplished using primers *nifH*1 (5’-ADNGCCATCATYTCNCC-3’) and *nifH*2 (5’-TGYGAYCCNAARGCNGA-3’) (^25^).

#### nifH bioinformatics

Paired-end sequencing reads were quality-trimmed (Phred score >20) and filtered to remove reads with more than 2 and 4 expected errors in the forward and reverse sequences, respectively. The DADA2 pipeline was used for sample inference, followed by paired-reads merging and sequence table construction. Taxonomic annotation was performed using mothur’s implementation of the naïve Bayesian classifier (^26^) with the *nifH* ARB reference database described by Schloss et al. (^27^). Additional information is provided in the supporting information. Diversity metrics—including richness, evenness, and Shannon index—were calculated to assess diazotroph alpha diversity across environmental conditions. Beta-diversity was inferred using Bray-Curtis dissimilarity and visualized using principal coordinate analysis (PCoA) to account for the effect of environmental conditions on community structure. Relative abundance of diazotrophic phyla was calculated and compared across lifestyles and seasons to identify key community members and their distribution patterns.

#### Chemical analysis of porewater and sediment

Organic measurements were performed in a TOC analyzer (Analytik-Jena Multi N/C 3100, Germany, detection limit = 0.3 mg L^-1^). Inorganic nitrogen species (NO ^-^, NO ^-^, NH ^+^) and phosphate (PO_4_^3-^) concentrations were measured by a flow injection autoanalyzer (Lachat Instruments QuikChem 8000, detection limit = 0.05 and 0.03 μmol L^-1^ for dissolved inorganic nitrogen, DIN, and PO_4_^-^, respectively).

#### Statistical Analyses

Diazotroph abundance, N_2_ fixation rates, and N_2_ fixation rates per-cell were quantified across two microbial lifestyles (biofilm-associated and free-living), two redox conditions (aerobic and suboxic), and two seasons (summer and winter). Number of replicates (n) per variable was between 2—7. More statistical details are provided in the supporting information.

## Results and Discussion

### Diazotroph abundance and N_2_ fixation rates in the hyporheic zone

Regardless of the oxic conditions and their lifestyles (i.e. free-living or biofilm-associated cells), diazotrophs comprised ≤9.6 % of the total bacterial abundance (Figure 1A–D, S2). It was assumed that detected cells after immunolabeling the nitrogenase enzyme have actively fixed N_2_ (i.e., diazotrophs) and differed from total bacteria (stained only by SYBR green). Abundance of free-living diazotrophs in the aerobic zone ranged from 4.4 to 47 × 10^6^ cells L^-1^ and 0.05 to 5.1 × 10^6^ cells g^-1^ for those associated with biofilms (Figure 1A, B). In contrast, in the suboxic zone, the free-living diazotrophs ranged from 9.3 to 770 × 10^6^ cells L^-1^, while biofilm-associated diazotrophs ranged from 0.03 to 0.9 × 10^6^ cells g^-1^.

**Figure 1.**
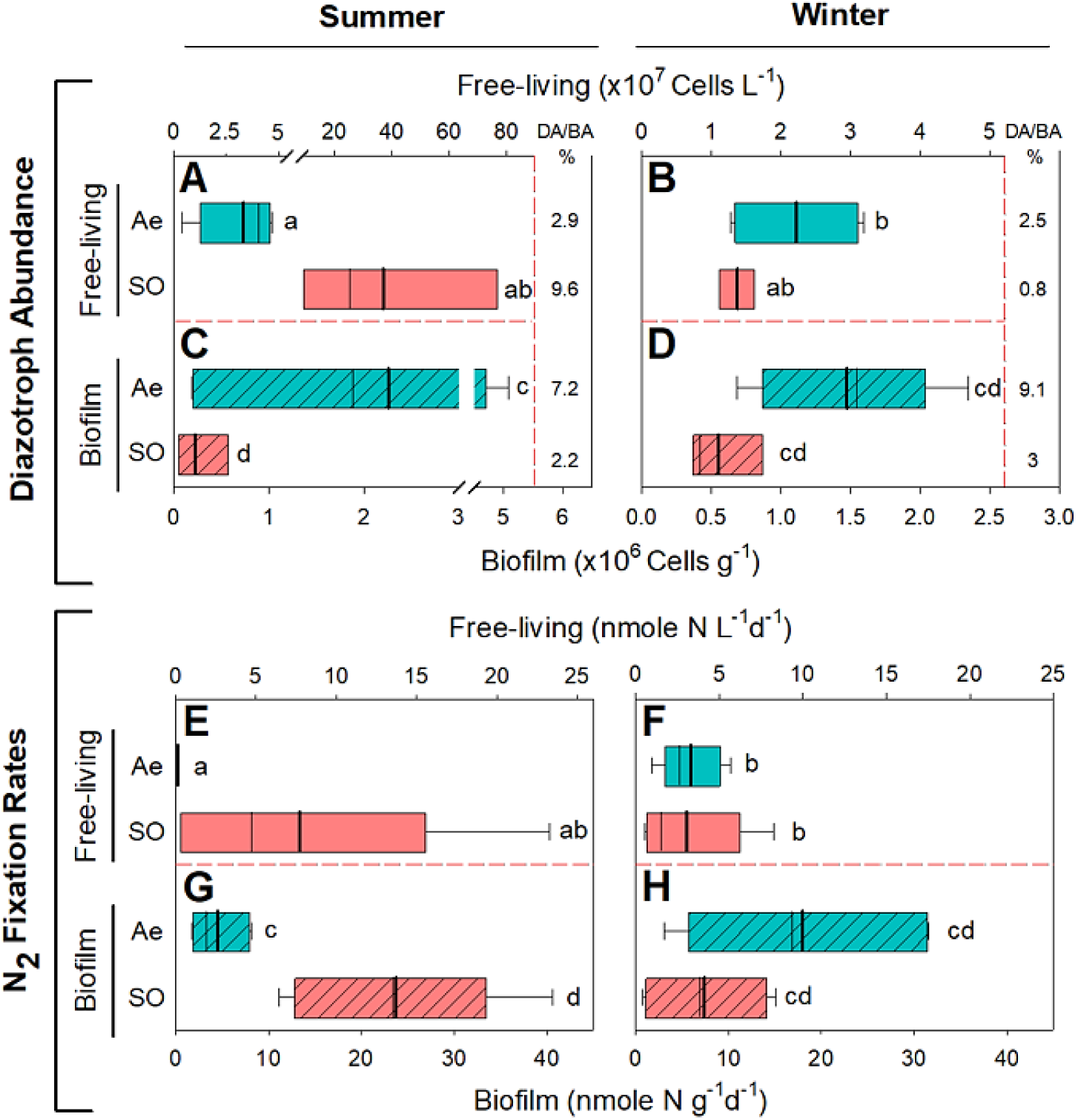
Diazotroph and bacterial abundance (A-D) and N_2_ fixation rates (E-F) sampled from the hyporheic zone during summer (A, C, E, G) and winter (B, D, F, H). Top x-axis for all panels corresponds to the free-living lifestyle and the bottom x-axis corresponds to biofilm-associated fractions for all plots. The y-axis for all plots reflects redox zonation and lifestyle. Aerobic (cyan), Ae, and suboxic (red), SO, are each plotted for both lifestyles, free-living (pastel colors) and biofilm (diagonal hatching). % BA indicates the compositional percentage of diazotrophs from the total bacterial community abundance (A-D). Box plots (A-H) portray interquartile range (25^th^ and 75^th^ percentiles) with whiskers extending to the 5^th^ and 95^th^ percentiles with a mean line (thick) and a median line (thin). Letters represent significant differences *p <* 0.05 determined by ANOVA and ART ANOVA analysis. Post hoc analysis was performed using Tukey’s test. ART ANOVA revealed a significant DO × Season interaction in both free-living and biofilm-associated lifestyles (FL *p =* 0.047, η² = 0.26, Bio *p =* 0.0029, η² = 0.40—0.45).

The abundance of free-living diazotrophs was highly affected by seasonality, decreasing by 1.7 times in the aerobic zone and by 32 times in the suboxic zone from summer to winter (Figure 1A, B). Nonetheless, during the summer, the average number of free-living diazotrophs quantified in the aerobic zone was 10 times lower (37 ± 22 × 10^6^ cells L^-1^) than those measured in the suboxic zone (Figure 1A). In contrast, during the winter, the number of free-living diazotrophs (22 ± 9 × 10^6^ cells L^-1^) was 2 times higher in the aerobic zone than the suboxic zone (11 ± 3 × 10^6^) (Figure 1B). Contrarily, the abundance of diazotrophs associated with biofilms was consistently higher in the aerobic than suboxic zone: by 17 times during the summer and 3.2 times during the winter (Figure 1C, D). Altogether, it was evident that oxic conditions and seasonality led to significant changes in the abundance of free-living diazotrophs (F_1,9_ = 5.55 and 7.91, *p =* 0.043 and 0.02, respectively), with higher abundances in suboxic zones and summer seasons. Diazotrophs associated with biofilms were mostly affected by oxic conditions, regardless of season (ANOVA, F_1,13_ = 11.6, p = 0.005), with higher abundance in aerobic than under suboxic conditions.

N_2_ fixation rates by free-living diazotrophs were barely detectable at the aerobic zone during the summer, increasing to 3.3 ± 1.9 nmol N L^-1^ d^-1^ in the winter (Figure 1E, F). N_2_ fixation rates at the suboxic zone by free-living diazotrophs ranged widely (0.3 to 23 nmol N L^-1^ d^-1^) yet did not differ greatly by season. However, N_2_ fixation rates by diazotrophs associated with biofilms found in the aerobic zone during the summer (4.6 ± 3.1 nmol N g^-1^ d^-1^) were 4 times lower than those measured during the winter (18 ± 12 nmol N g^-1^ d^-1^) (Figure 1G, H). Inversely, in the suboxic zone, these rates were ∼3 times higher in the summer, compared to the winter.

These results indicate that both DO and season significantly affected N_2_ fixation rates across diazotroph lifestyles (FL: *p =* 0.047, η² =0.26; Bio: *p =* 0.0029, η² =0.40). This interaction was driven by higher suboxic N_2_ fixation during summer compared to winter, contrasting with comparably stable aerobic rates. Delving into the disparity in abiotic conditions between aerobic and suboxic zones during summer highlights the central role of DO in modulating diazotroph abundance and N_2_ fixation rates along the streambed depth profile. Compared to the aerobic zone in summer, the suboxic zone was characterized by lower DO concentrations and N:P ratios below the Redfield ratio (Table S1). These conditions likely spur diazotroph abundance and corresponding N_2_ fixation rates, consistent with patterns reported in marine environments (^1,28^). During winter, reduced availability of organic matter, extremely high N:P ratios, and higher DO saturation, even at depth (∼16 cm, up to 40 %), led to a reduction in the numbers of free-living diazotrophs.

Diazotrophs associated with biofilms were expected to be less constrained by the porewater conditions due to the collective metabolic activity of the bacterial consortia and the presence of extracellular polymeric substances (EPS) that impair diffusion-based processes and promote the formation of suboxic microzones (^29–31^). Unique conditions can be found within biofilms that possibly spur diazotrophy, including minimal DO micro-zones (^15,29,32,33^) and specific loci with abundant labile carbon and reduced concentrations of DIN due to enhanced enzymatic processes (^12,34^). Overall, the activity of the bacterial consortia associated with these diazotrophs could supply beneficial metabolites while consuming both ambient DO and available N, thus increasing the demand for fixed N_2_ (^15,29,35–37^). Still, N_2_ fixation rates were often higher in the suboxic zone due to reduced competition of DO with N_2_ over the catalytic site of the nitrogenase enzyme (^15^).

N_2_ fixation rates per-cell were determined to proffer a direct comparison between diazotrophs at different locations and lifestyles. Notably, these per-cell trends diverged from patterns observed in diazotroph abundance, indicating that DO availability and seasonality modulate diazotroph activity independently of population size. Overall, average N_2_ fixation rates per single biofilm-associated cell were between 6—>72,000 times greater than those measured by free-living diazotrophs, with a 377-fold average increase (Figure 2). During summer, N_2_ fixation rates by a free-living cell at the aerobic zone were below the detection level, while the corresponding rates by those associated with biofilms were 15 ± 14 femtomole N cell^-1^ d^-1^ (Figure 2A). Although a similar trend was observed during winter, N_2_ fixation rates per-cell were detectable in aerobic free-living cells (0.13 femtomole N cell^-1^ d^-1^) and 1.4 times higher in biofilms (Figure 2B).

**Figure 2.**
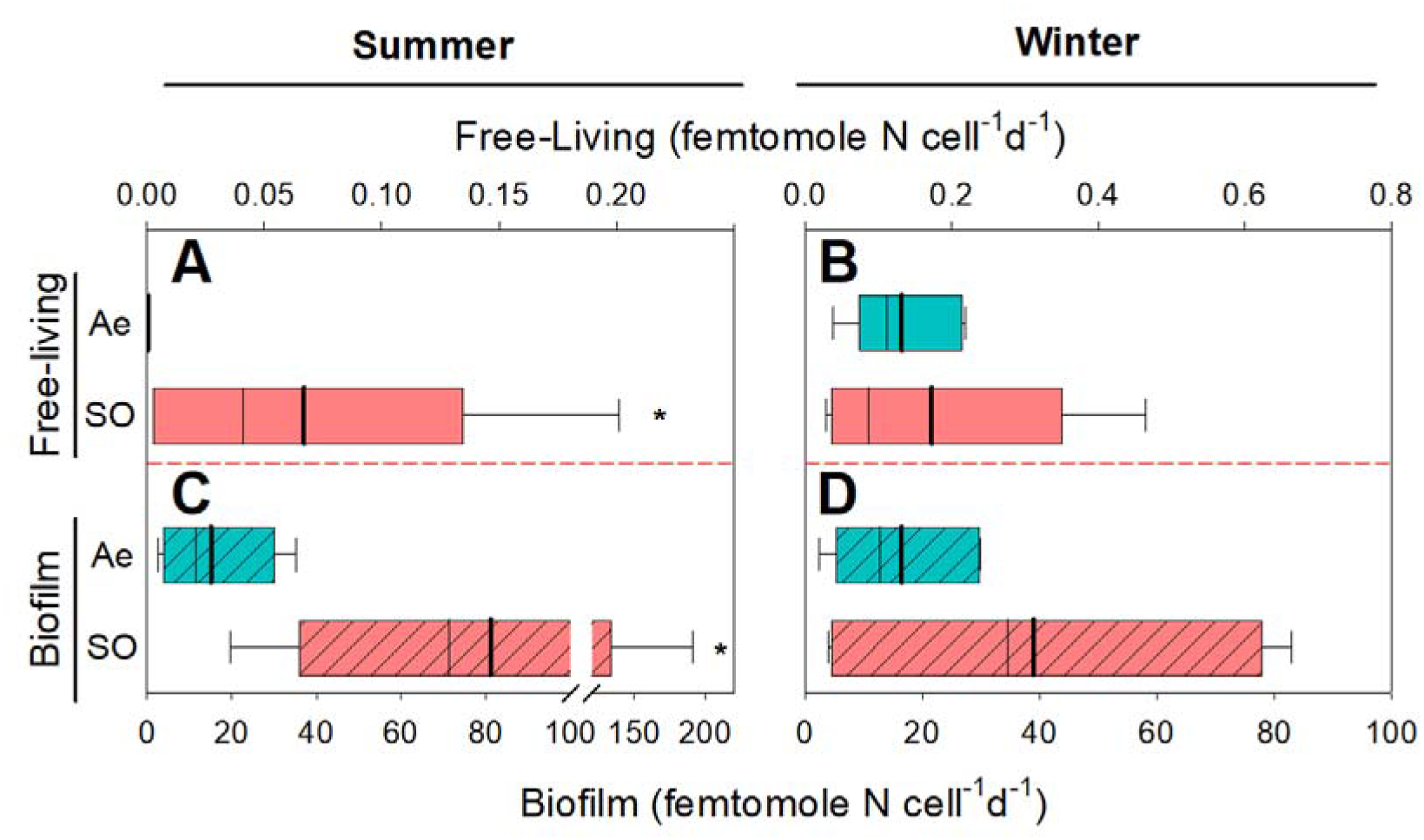
N_2_ fixation per-cell is calculated by dividing individual response measurements of N_2_ fixation rates by corresponding diazotroph cell numbers for free-living (A, B) and biofilm-associated (C, D) diazotrophs. These values were determined for diazotrophs sampled during the summer (A, C) and winter (B, D) under aerobic (Ae, cyan), and suboxic (SO, red) conditions. Top x-axis corresponds to the free-living fraction, and the bottom to the biofilm-associated (diagonal hatching) fraction. Box plots (A-D) portray interquartile range (25^th^ and 75^th^ percentiles) with whiskers extending to the 5^th^ and 95^th^ percentiles with a mean line (thick) and a median line (thin). Letters represent significance *p <* 0.05 determined by ANOVA and ART ANOVA analysis. Post hoc analysis was performed using Tukey’s test.

Irrespective of the redox zonation in summer, N_2_ fixation by a single cell associated with biofilms was significantly higher (by 1545 times, F_1,37_ = 43.224, *p <* 0.001) than the free-living fraction and ranged between 2.6 and 187 femtomole N cell^-1^ d^-1^ (Figure 2C). During winter, N_2_ fixation per-cell associated with biofilms increased from 16 ± 12 femtomole N cell^-1^ d^-1^ in the aerobic zone to 39 ± 40 femtomole N cell^-1^ d^-1^ in the suboxic zone (Figure 2B, D). Overall, these rates were 166 times greater than those measured for a free-living diazotroph cell.

Nonetheless, free-living diazotrophs in winter fixed N_2_ at significantly higher rates per-cell than in summer (*p <* 0.01). Biofilm-associated cells fixed N_2_ at comparable rates in the aerobic zone across seasons, whereas rates in the suboxic zone were over two times higher during summer than during winter (Figure 2C, D). Suboxic biofilm-associated diazotrophs fixed significantly higher N_2_ per-cell than aerobic diazotrophs (*p =* 0.009, η² ≈ 0.32). In free-living diazotrophs, significantly higher fixation per-cell was measured in winter (*p =* 0.009, η² ≈ 0.31).

Quantifying N_2_ fixation rates per-cell illumines a striking disparity in diazotrophic activity between microbial lifestyles in the hyporheic zone. Previous reports indicate that DO, availability of organic matter, and the stochiometric ratios of C:N:P holds a central role in governing diazotrophy (^7,8,28^). However, it is evident that regardless of their lifestyle, the greatest driver of N_2_ fixation per-cell within this streambed environment was DO saturation. Irrespective of seasonal variability, the highest N_2_ fixation per-cell was attributed to diazotrophs associated with biofilms. As previously discussed, biofilms (similarly to planktonic aggregates) often provide conditions that support N_2_ fixation even in environments that could otherwise be considered adverse to diazotrophy (^38–40^).

### Microlocalization of diazotrophs associated with biofilms in the hyporheic zone

Diazotrophs that synthesized the nitrogenase enzyme were found to reside in a tight nexus with bacteria as part of greater biofilm complexes (Figure 3). Biofilms located in the aerobic zone were observed to be consistently enveloped by thick EPS—several tens of microns (Figure 3A, B). Active diazotrophs associated with these biofilms were embedded within that EPS matrix and often congregated in small clusters adjacent to neighboring bacteria (Figure 3). Biofilms collected from the suboxic zone appeared to have only a thin EPS zone (<20 µm) with sparsely dispersed polysaccharide globules (Figure 3C, D). Similarly to the aerobic zone, total bacteria and diazotroph cells attached to the sediment particles were also captured in a tight nexus. Nonetheless, these diazotroph cells were often directly exposed to suboxic porewater. The thin EPS zone found at deep riverbed layers under suboxic conditions was possibly the result of heterotrophic degradation of polysaccharides and proteins that comprise EPS over time (^41,42^). As previously reported, increased production of EPS can occur due to high nutrient loads (^43,44^). A recent study on nitrogen-fixing *Paenibacillus polymyxa* reported that EPS production was intensified under aerobic conditions as a stress response to DO (^45^). This thick EPS envelope can lead to steep oxic gradients within the biofilm due to bacterial respiration and lower diffusion rates, forming localized oxygen minimum microzones (^29,32,33^) that minimize damage to the nitrogenase enzyme.

**Figure 3.**
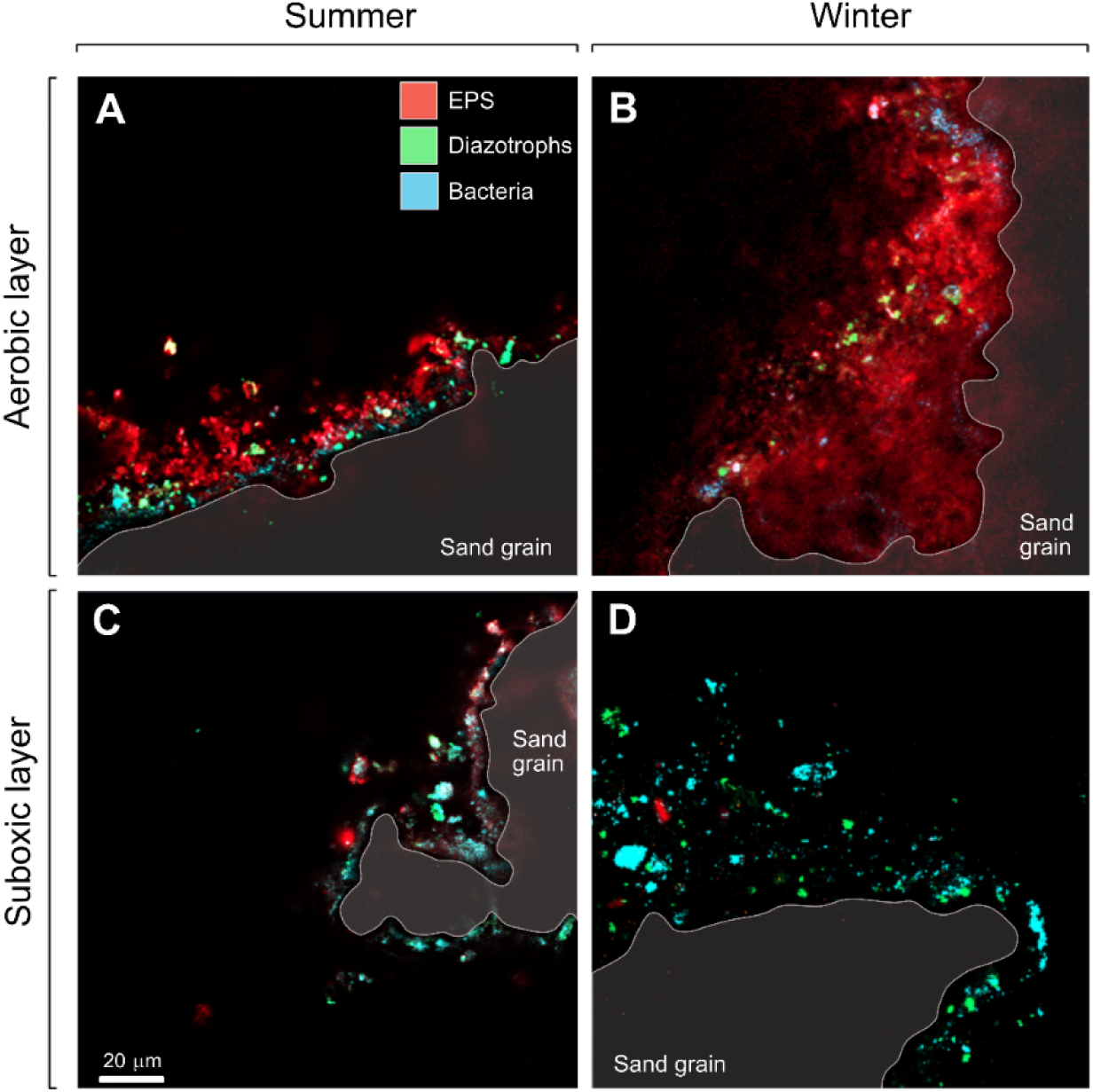
Confocal laser scanning micrographs of biofilms associated with sand grains collected from the hyporheic zone at the Jordan River during the summer and winter (left and right panels, respectively). These biofilms were attached to the sand in both aerobic (A, B) and suboxic layers (C, D). The sediment grain was marked by a grey contour line and semi-transparent shading. Extracellular polymeric substances (EPS) were stained with Concanavalin A, which binds to the polysaccharide matrix (red). Total bacteria were stained by DAPI (cyan), while diazotrophs were distinguished by immunolabeling the nitrogenase enzyme (green).

The C:N ratio of the polysaccharide matrix that comprises the EPS is often higher than the Redfield ratio (>6.6:1) due to the rapid degradation of nitrogen molecules compared to carbon (^46^). These conditions, including suboxic microzones, can support the growth and activity of diazotrophs that colonize biofilms despite conditions in the surrounding environment that inhibit N_2_ fixation (^15,29^). Due to the consistently low saturation of DO (<39 %) at suboxic layers, it is expected that diazotrophs associated with biofilms will gain distinct metabolic advantages that catalyze diazotrophy. Such advantages may derive from enhanced accessibility to labile organic matter, supplied through the biodegradation process by the heterotrophic community that comprises these biofilms (^13,47^). These micrographs highlight the intimate spatial environment in which diazotrophs cohabitate with the bacterial consortiums and suggest functional interactions.

### Diversity of diazotrophs identified in the hyporheic zone

PCoA analysis of the diazotroph community in the hyporheic zone revealed clear clustering by lifestyle, namely as free-living and biofilm-associated cells (Figure 4A). Free-living diazotrophs were observed to cluster in aerobic and suboxic layers (yet not statistically significant; *F =* 5.56, *p =* 0.10), indicating that DO availability governs these communities. Seasonality did not affect beta diversity, suggesting that lifestyle and redox conditions are the primary drivers of community composition. Although we found little disparity between the effects of DO availability and seasonality on alpha diversity, there was a large effect size (Cohen’s *d* = 1.12) on diversity by lifestyle. A large effect size, computed by Cohen’s *d*, suggests meaningful trends regardless of the insufficient statistical power of the test to prove significance (Figure 4B). Biofilm-associated communities exhibited higher Shannon diversity than free-living communities, 3.7 and 3.3, respectively (*t* (7.97) = –1.93, *p =* 0.09, *d* = –1.12).

**Figure 4.**
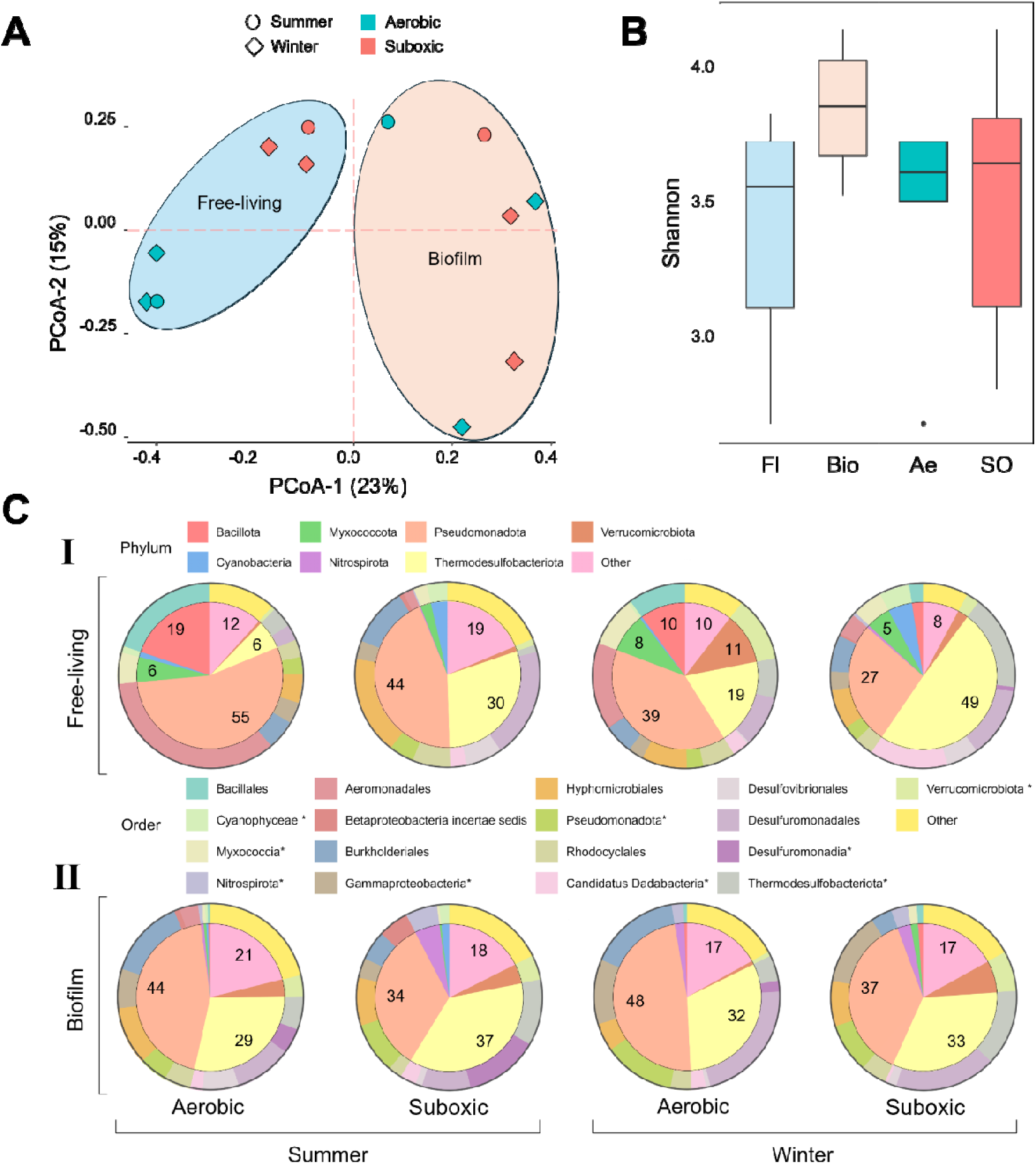
Principal coordinate analysis (PCoA) showing distance in diazotroph diversity derived from *nifH* analyses categorized by DO gradient (cyan or red), season (Summer – circle, Winter –diamond), and lifestyle (A). Diversity as a response to season (summer, winter), redox zone (Aerobic – Ae, Suboxic – SO), and lifestyle (Free-living – FL, Biofilm-associated – Bio), from left to right on the Shannon index (B). Phylum (inner pie chart) and Order (outer pie chart) for dominant taxa within free-living (I) and biofilm-associated (II) diazotrophs in the Summer (left) and Winter (Right) (C). Only taxa with 5 % or greater representation in any one sample are named, while taxa below this threshold are labeled “Other”. Asterisks (*) indicate that the specific taxa are unclassified. All taxa can be found in supplementary information. Welch’s t-tests were performed on B in addition to two-way ANOVA (Lifestyle×DO) and three-way ANOVA (Lifestyle×DO×Season) with Cohen’s d to test for effect size in a 95 % CI. PERMANOVA with pairwise comparisons were run on A with 999—9999 permutations and adjusted with Benjamini-Hochberg.

Interestingly, the alpha diversity varied more at suboxic depths than in the aerobic zone, with a standard deviation roughly two times higher and a wider IQR (SD = 14, CV = 18 and SD = 7.6, CV = 9.5, respectively) (Figure 4B). The Shannon diversity during the winter was less variable than in summer, with a coefficient of variance of 11 % as compared to summer’s 16 %. We deduced that the diazotroph community structure in this riverbed is modulated by lifestyle (Figure 4A, B), as previously reported for other types of bacterial populations in subsurface environments (^48^).

Free-living diazotrophs such as unicellular cyanobacteria and NCDs are more susceptible to oxic conditions in the surrounding environment than those associated with biofilms (^19,49^). Biofilms can regulate internal redox conditions and maintain buffers within themselves from the surrounding environment, which mitigates the impact of oxygen availability and the vacillations in abiotic conditions associated with seasonality (^19,29,48,50^). Therefore, it was not surprising that diazotrophs found in the porewater under varying oxic conditions formed segregated community clusters.

Overall, Pseudomonadota dominated the diazotroph community found in the hyporheic zone, comprising 27—55 % of the population, reaching the highest relative abundances under aerobic conditions (Figure 4C, S3, Table S3). This phylum is ubiquitously present in aquatic environments, often found as facultative or obligate anaerobes (^51,52^). Within this phylum, Gammaproteobacteria, Betaproteobacteria, and Alphaproteobacteria were dominant in descending order of relative abundance (16, 14, and 12 %, respectively), all commonly identified as aquatic NCDs (^53,54^). It is suggested that metabolic plasticity (i.e., heterotrophy, chemotrophy, or mixotrophy) enabled these diazotroph orders to thrive in both porewater and biofilms under varying oxic conditions. A similar trend corresponded with Thermodesulfobacteriota, comprising 6—49 % of the diazotroph phyla (Figure 4C). This phylum comprises mostly anaerobic, gram-negative bacteria, often dominated by various sulfate-reducing bacteria such as identified herein: Desulfovibrionia, Desulfuromonadia, and Candidatus Dadabacteria. The ubiquitous presence observed of *Geobacter, Desulfuromonadales*, and *Desulfovibrio* indicates the co-occurrence of Fe(III) and sulfate reduction alongside N_2_ fixation, previously shown to be processes that are coupled together (^55–57^). Although *Geobacter* and *Oceanidesulfovibrio* are strict anaerobes, they were found in all oxic layers, intimating the possible community links between the hyporheic layers. Such links may arise from advective porewater exchange and vertical transport of cells and metabolites from adjacent sediment layers (^58,59^).

Free-living communities included higher proportions of *Paenibacillus*, *Tolumonas*, Cyanophyceae, and Myxoccocia (8, 14, 3, and 6 %, respectively) than biofilm communities (<1 %) (Figure 4C I, II). Free-living diazotrophs often comprise Bacillota, Myxococcota, and Cyanobacteria phyla including the corresponding taxa Bacillales *Paenibacillus*, an unclassified Myxococcia, an unclassified Cyanophyceae, Aeromonadales *Tolumonas* and Cyanophyceae (Figure 4C I, Table S3). Among these free-living diazotrophs, members of *Paenibacillus* likely contribute heavily to N_2_fixation, often being facultative anaerobes with broad fermentative metabolic capacities, with 20 species confirmed to fix N_2_ (^60,61^). Myxococcia is a predominantly aerobic bacteria (^62^), so preferentially inhabits free-living fractions where diffusion of oxygen is more linear and pervasive compared to biofilms. Cyanophyceae commonly display mixotrophic behavior and the capacity for N_2_ fixation, enabling habitation in low-light conditions (^63,64^). Interestingly, more Cyanophyceae were identified in the aphotic suboxic layers than in the aerobic (3 % in suboxic and 0.5 % in aerobic), further accentuating their potential mixotrophic capacity (Figure S3).

Biofilm-associated communities exhibited high relative abundances of Thermodesulfobacteriota (33 %), predominantly represented by *Geobacter* (14 %), alongside additional lineages including *Desulfuromonadia, Desulfovibrio, Oceanidesulfovibrio,* and *Desulfobulbus*. These taxa are well-established contributors to suboxic electron transfer, sulfur and metal cycling, as well as N_2_ fixation (^21,57,65^). *Desulfovibrio* and *Desulfobulbus* are known to be aerotolerant (^65^). Biofilms in aerobic environments comprised high proportions of Desulfuromonadia, indicating the potential for sustaining suboxic niches within these matrices (Figure 4C II). *Geobacter* is a well-studied N_2_ fixer and has the propensity to form biofilms (^64,66^). Winter aerobic biofilms were notably enriched in *Geobacter* (22 %), likely due to its capacity to stratify itself within biofilms and colonize the innermost layers (^67^).

Altogether, diazotroph communities in the hyporheic zone were structured by interactions between lifestyle and DO and were less influenced by seasonality. While biofilm assemblages remained stable through oxic vacillations and even harbored strict anaerobes under oxygenated conditions, free-living diazotrophs were more susceptible to the changes between aerobic and suboxic layers and differed sharply. These patterns exemplify the importance of lifestyle in modulating how diazotrophs respond to redox zonation, as evinced by the ubiquity of metabolically versatile groups such as *Pseudomonadota* and *Geobacter* to persist across different environments.

### Comparing diazotrophy in the hyporheic zone across freshwater environments

Comparing diazotroph activity within the hyporheic zone (free-living cells and biofilm-associated) to the overlying water column of the Jordan River was achieved by converting measured N_2_ fixation rates into cubic meters as a converging unit (detailed in the supplementary methods, Table S1). N_2_ fixation rates by biofilm-associated diazotrophs (3.6 × 10^7^ nmol N m^-3^ d^-1^) were 5 × 10^3^-fold greater than free-living cells found in the porewater (Table S5). Compiling the above, N_2_ fixation rates within the hyporheic zone by free-living cells and those ascribed to biofilms were 1.5 × 10^5^-fold higher (3—4 × 10^7^ nmol N m^-3^ d^-1^) than the rates measured in the overlying water column (237 nmol N m^-3^ d^-1^, ^6^) from the same sample location (Table S5). The N_2_ fixation rates measured within this hyporheic zone were 2.8 × 10^2^-fold greater than rates reported from the water columns of other rivers and 5.8 × 10^2^-fold larger than freshwater lakes reported at the global scale (^68^). Moreover, the rates measured herein were nearly five-fold greater than rates reported for inland sediments (^68^), further highlighting the potential role of subsurface diazotrophy as a new N source.

The pronounced disparity between N_2_ fixation rates in the hyporheic zone and the overlying water column indicates that this streambed environment is a biogeochemical hotspot for diazotrophs. These results and corresponding insights add to the already developed knowledge that the hyporheic zones act as a natural bioreactor, facilitating a wide range of biogeochemical processes (^20,69,70^). It is likely that a large fraction of this newly fixed N is rapidly assimilated into microbial biomass or retained through adsorption to sediments and EPS in biofilms (^71^). Due to hydrological connectivity, the intimate nexus of flora and diazotrophs at such fluvial ecosystems—especially at the hyporheic zone—produces interconnected physicochemical processes (^72–74^). Therefore, it is suggested that this subsurface hotspot for diazotroph activity and the dynamic hydrologic exchange can support the exchange of fixed N from the hyporheic zone into the overlying stream (^71,75^).

Moreso, hydrological connectivity can further transport this new fixed nitrogen to downstream ecosystems (^72–74^). Deductions from these results along with recent global syntheses intimate that inland waters and freshwater subsurface may contribute to global N_2_ fixation more than previously thought (^68^).

## Conclusion

This research provides the first comprehensive insights into streambed diazotrophy. It was found that streambed diazotrophs made up ≤9.6 % of total bacteria in the hyporheic zone and were consistently more enriched in biofilms. Biofilm-associated diazotrophs were buffered from surrounding conditions, despite immense differences in DO and nutrient stoichiometry, maintaining higher N_2_ fixation rates per-cell and rich diversity. Conversely, free-living diazotrophs found in the porewater diverged sharply between N_2_ fixation rates and population diversity across different redox zonations. Per-cell N_2_ fixation in biofilms reached up to 377-fold higher than free-living cells. Sequencing the *nifH* indicated that the community structure of these streambed diazotrophs was shaped foremost by lifestyle and DO, while only marginally by seasonality. The results and corresponding insights highlight that the hyporheic zone is a diazotrophy hotspot and further shift the paradigm related to the roles of N_2_ fixers along the freshwater continuum.

## Acknowledgements

We thank Dr. Eyal Geisler for his assistance with sample analysis and the ranger Noam Revach for technical support. This study was supported by the Israeli Science Foundation (grant number 944\21 to E.B-Z).

## Author Contributions

RL and SA and EB-Z conceived and designed the sampling campaigns. RL and HS measured the N_2_ fixation rates together with EG. RL and CV analyzed all the bioinformatic data. The manuscript was written mainly by RL and EB-Z, whereas all authors commented and contributed to previous versions of the manuscript. All authors read and approved of the final manuscript.

