## Supplementary material for "Lifestyle modulates riverbed diazotrophy: from abundance and activity to diversity of free-living and biofilm-associated N_2_ fixers": SI: 20260415_Lander et al_SI.docx

* Corresponding author:

**Materials and Methods**

**Sampling strategy**

Sediment samples and porewater were collected from the Jordan River at a downstream site (Gesher Arik, 32º 5408.29N; 35º 3651.12E) (Figure S1).

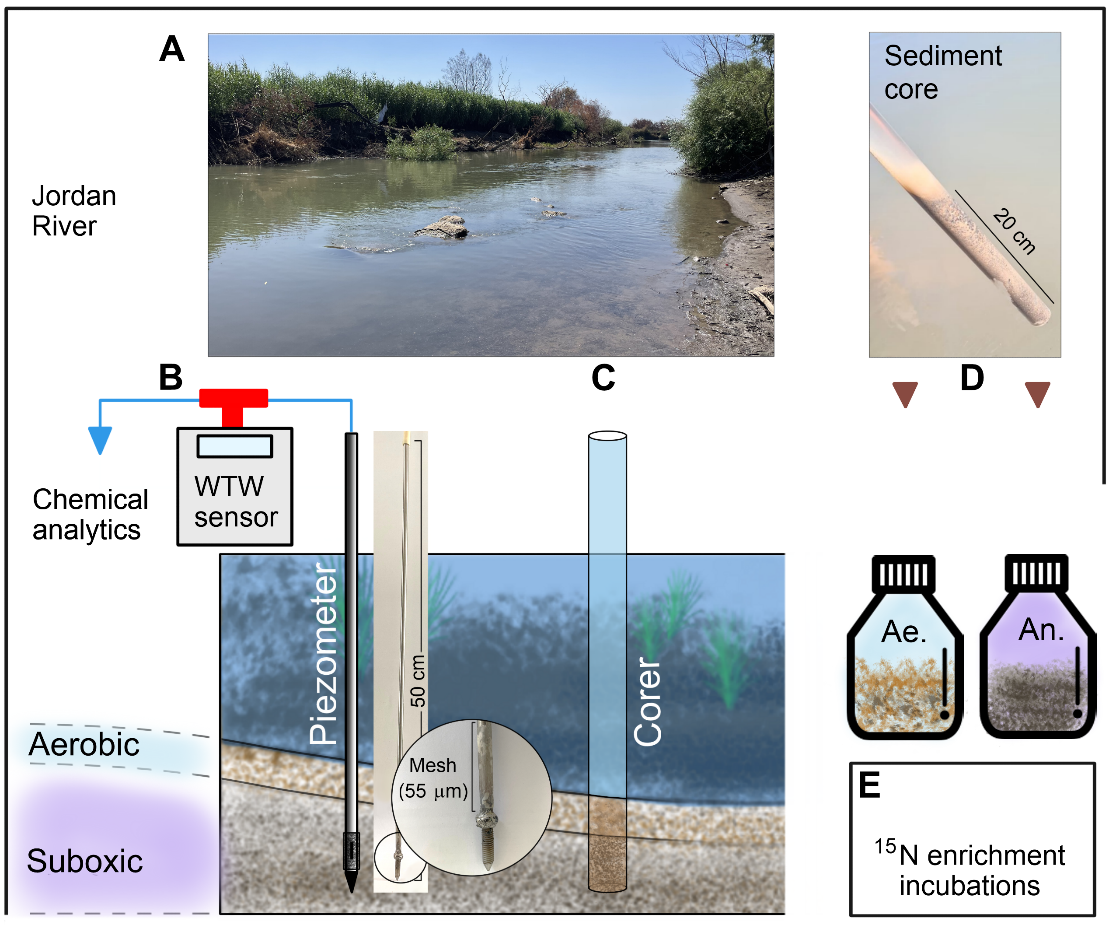

**Figure S1.** Sampling design and on-site workflow. Downstream view of the sampling site at the Jordan River (A). Schematic and corresponding pictures of the piezometer used to sample the hyporheic zone, highlighting the aerobic and suboxic zones (B). Corer schematics (C) and a representative core collected from the hyporheic zone (D). Illustration of the core-microcosms with sediment and porewater for aerobic and suboxic zones, which were incubated with ^15^N_2_ (E).

*Nitrogenase Immunolabeling*

Samples were centrifuged (4,000 rpm, 10 min), washed three times with PBST (0.1 % Triton X-100 in phosphate-buffered saline, pH 7.2, Sigma Aldrich, ^1^), and incubated with a primary anti-*nifH* antibody (3 μg mL^-1^, Agrisera-AS01 021A, Sweden) in PBST with bovine serum albumin (1 mg mL^-1^, BSA, Sigma Aldrich A2153). After 1 h incubation, samples were centrifuged and washed again, labeled with a secondary antibody solution (6 μL/mL, goat anti-chicken IgY conjugated to fluorophore; Ex: 495 nm, Em: 519 nm) in PBST with BSA (1 mg/mL), followed by a 45 min incubation. Labeled samples were washed with 0.4 % NaCl (3 times) to remove unbound antibodies. All samples were labeled alongside their corresponding sterilized samples as well as *Vibrio* *natriegens* as the positive control. Subsamples (250 μl) were analyzed in the acoustic focusing flow cytometer and directly compared to the sterile replicates. Negative controls include 0.4 % NaCl with the secondary antibody to check for unspecific binding, *E. coli* with both primary and secondary antibodies, and *Vibrio natriegens* with the secondary antibody. Additionally, a subsample was aliquoted from each sample and autoclaved, immunolabeled, quantified, then subtracted from the actual sample quantity due to background noise from clay particles.

Total bacterial abundance was determined by staining a subsample with 0.5 nM SYBR Green I (S7563, Invitrogen, final concentration, 1 nM) for 15 min in the dark. Stained samples were then quantified by a flow cytometer (Applied Biosystems), using an excitation wavelength of 450 nm and emission detector of 520 ± 30 nm. Phototrophic microbes were quantified by the detection of the autofluorescence of chlorophyll a and phycoerythrin with an emission detector of 574 ± 26 nm (^2^). Beads (0.93 μm) were used as a standard to compare size fractions. Negative controls included autoclaved samples that were stained in the same manner as the above. Note, it was assumed that detected cells after immunolabeling the nitrogenase enzyme have actively fixed N_2_ (i.e., diazotrophs) and differed from total bacteria (stained only by SYBR green).

*Preparation of ^15^N_2_ enriched water*

The dissolved stock of ^15^N_2_ was prepared according to Geisler et al. 2023. MilliQ water was added to a gas tight bottle (750 mL) and degassed using under vacuum (15 psi) with a dedicated membrane (G543, MiniModule) and custom-made tubing for 24 h at 30ºC. The degassed water bottle was injected with 15 mL of ^15^N_2_ gas (99%, Cambridge Isotopes, Lot # NLM-363-PK) at a ratio of 1.5% v:v through dedicated tubing. The enriched bottle was placed on a shaker in a cold room (4 °C) for three days, until the gas bubble completely dissolved.

*EA-IRMS quality control and calibration*

Dry samples were packed into tin capsules and analyzed for N quantification and δ^15^N measurements of both natural abundance and enriched samples, using an elemental analyzer (EA; Flash 2000 HT, Thermo Scientific, Milan, Italy) coupled with an isotopic ratio mass spectrometer (IRMS; Delta V Plus, Thermo Scientific, Bremen, Germany). Secondary standards were used for isotope calibration and quantification. Standards spanning the isotopic range of the samples bracketed the measurements to ensure accuracy, while standards with varying nitrogen amounts were used for quantitative calibration. A repeated reference standard was analyzed throughout each analytical sequence to monitor instrumental stability and correct for potential drift.

A set of secondary standards was used to bracket and ensure the accuracy of the isotope analysis, for quantification, and to correct for potential drift. The analytical procedures followed the approach described in detail in Geisler et al. (^3^). Briefly: Quartz crucibles were used and replaced every 10–20 samples to prevent clogging or incomplete combustion in the reactor. The accuracy of the isotopic measurements was verified by analyzing international reference materials at the beginning and end of each analytical sequence. The following reference materials were used: caffeine (USGS62, δ¹⁵N = +20.17‰), glycine (USGS64, δ¹⁵N = +1.76‰), and L-glutamic acid (USGS40, δ¹⁵N = −4.52 ‰(. The standards bracketed the isotopic values of the samples, ensuring accuracy across the full range of observed values. USGS64 was analyzed every 4–5 samples throughout each sequence to monitor analytical precision (< ±0.3‰) and instrumental stability and to correct for potential drift. Nitrogen quantification was performed using a calibration curve based on peak amplitudes of the reference standards. The calibration was linear across the measured range (R² > 0.99). Blanks were measured at the beginning of each sequence and were routinely subtracted. Blank contributions were negligible relative to sample signals.

*DNA extraction and nifH Amplification*

*nifH* genes were amplified with primers (Hylabs, Israel) using Takara Taq (R011, Takara, Japan) in a polymerase chain reaction (PCR, Life Eco, Bioer Technology, China) with two cycles. The first cycle was run with reverse primer: TTYTAYGGNAARGGNGG and forward: ATRTTRTTNGCNGCRTA. The second cycle was run with the reverse and forward primers ADNGCCATCATYTCNCC and TGYGAYCCNAARGCNGA, respectively. The first cycle was set to 94 °C for 5 min, cycled 30 times at 94 °C 1 min, 50 °C 1 min, 72 °C 1 min, and then extended at 72 °C 1 min. The second cycle was run at 94 °C 5 min, then there were 30 cycles of 94 °C 1 min, 57 °C 1 min, 72 °C 1 min, and an extension at 72 °C 1 min. Each PCR run was accompanied by a positive (*V. natriegens*) and negative (double-distilled water, DDW) control plus an additional negative control for the second cycle of DDW with PCR reagents after the first cycle. PCR products were all visualized using gel electrophoresis.

*Normalizing biofilm-associated N_2_ fixation rates to a volumetric unit*

The contribution of N_2_ fixation rates was estimated by converting the biofilm-associated N_2_ fixation rates (nmol N g^-1^ d^-1^) into a volumetric rate (nmol N m^-3^ d^-1^) (Table S4). Since each bottle contained multiple intact cores, the sediment was treated as a single, bulk core. First, all sediment samples in microcosms (100 mL) were weighed as wet mass, dried at 105 °C for 48 h, then weighed again for dry mass. The mass difference between wet and dry samples was attributed to porewater, and its volume was calculated assuming a water density of 1000 kg m⁻³. This allowed estimation of porosity as the ratio of porewater volume to bulk sample volume (100 mL), and the complementary solids fraction. Dry bulk density was determined from the dry mass divided by bulk sample volume, while grain density was calculated from the dry mass divided by the volume of solids (bulk volume minus porewater volume). Assuming the grains were spherical, the volume of a single particle was calculated from the assumption of a median diameter of 250 μm. From this, the total number of particles in a cubic meter of bulk sediment and the number of particles per gram dry sediment were determined. Particles per gram sediment were divided by measured N_2_ fixation rates to determine a per-particle N_2_ fixation rate and then scaled up to bulk cubic meter values by multiplying the per-particle rate by the number of particles per bulk cubic meter. This procedure allowed direct normalization of measured fixation rates to the bulk sediment scale (nmol N m^-3^ d^-1^) and an easy comparison by converting the already volumetric unit of the free-living rates.

*Chemical analysis of porewater and sediment*

Porewater and sediment samples were taken *in-situ* and frozen with 1 % 1 N HCl in acid washed VOC vials to arrest metabolic activity until analysis. Before analysis, sediment samples were mixed with EDTA and NaCl to make a slurry as described above, then sonicated to detach and suspend biofilms and their constituents. The slurry was allowed to settle for 5 min, then the supernatant was collected for further analysis alongside the porewater. Each sample vial was split into two fractions: total organic carbon (TOC) and dissolved organic carbon (DOC; following filtration through 0.45 μm filters). Samples were diluted to EC <1 mS cm^-1^ and to a total organic carbon, TOC, concentration of <100 ppm, then measured by a TOC analyzer (Analytik-Jena Multi N/C 3100, Germany, detection limit = 0.3 mg L^-1^). Inorganic nitrogen species (NO_3_^-^, NO_2_^-^, NH_4_^+^) and phosphate (PO_4_^3-^) concentrations were measured by a flow injection autoanalyzer (Lachat Instruments QuikChem 8000, detection limit = 0.05 and 0.03 μmol L^-1^ for dissolved inorganic nitrogen, DIN, and PO_4_^-^, respectively).

*Statistical Analyses*

Diazotroph abundance, activity, and per-cell activity were determined by three-way and two-way ANOVAs. Assumptions were tested by Shapiro-Wilk and Levene’s. Results were validated with ART and log(x+1) transformations to stabilize variances and reduce skewness. Effect sizes (η² partial) evaluated the magnitude of effects while Cohen’s d was used to quantify effect size between groups. When assumptions were violated, results were cross validated with the non-parametric Aligned Rank Transformation (ART) ANOVAs. Post-hoc comparisons were conducted using Tukey’s HSD for parametric models and Benjamini-Hochberg, BH-adjusted pairwise contrasts for ART models. Shannon diversity and richness were additionally analyzed with Welch’s t-tests. R (version 4.2) was used for all statistical analysis with packages: tidyverse, car, readxl, rstatix, broom, broom.mixed, officer, flextable, ARTool, emmeans, and effectsize, phyloseq, and vegan.

**Results**

**Physicochemical characteristics of the hyporheic zone**

The hyporheic zone in the downstream site of the Jordan River was highly dynamic and exhibited significant differences in porewater temperature and DO relative to the sampling depth (Table S1). The aerobic zone was characterized by DO concentrations of 78–89 % at depths of 1 cm during the summer. During winter, DO concentrations increased substantially (88—97 %), penetrating to depths of 11 ± 3.6 cm. This seasonal deepening of the aerobic zone corresponded with an 8 °C decrease in temperature, from 24 ± 1.3 °C in the summer to 16 ± 1 °C in winter. The decline in temperature, increasing DO solubility, and the lower oxygen demand by subsurface microbes during winter (^4,5^) resulted in a deeper aerobic zone compared to summer. Conversely, the suboxic zone during summer was characterized by low DO saturation (<4—20 %) at depths of 7 ± 6 cm and porewater temperatures of 30 ± 4.6 °C. It is suggested that during the summertime, leaf litter deposition and the senescence of phytoplankton blooms (^3,6,7^) enhanced microbial activity in the subsurface further reducing DO (^8^). In contrast, the suboxic zone during the winter deepened to 14 ± 4 cm with a DO saturation of 13—39 %, where porewater temperatures averaged 17.8 ± 1.5 °C.

The N:P ratio (23—394) at the aerobic zone was highly variable, nonetheless, much higher than the Redfield ratio (16:1, ^9^) regardless of the sampled season (Table S1). This N:P ratio most likely suggests a phosphorus deficiency as it is preferentially mineralized over nitrogen (^10,11^). However, at the suboxic zones the N:P ratio was at the same scale as the Redfield ratio during the summertime (8.7 ± 9.4), yet similar to the aerobic zone during the wintertime (263 ± 106). As compared to the overlying water, N:P ratio did not differ significantly except within the summer suboxic zone, with an N:P of nearly 10 times lower than the stream (84 ± 5) (^3^). C:N ratios were at lower or similar values (1.2—5.7) than the Redfield ratio (6.6:1), with the exception of the winter suboxic zone (8.6 ± 6.7) (Table S1). It is likely that these low C:N ratios derived from high inputs of nitrogen compounds (mostly nitrate, 65—194 µmol L^-1^) from the overlying water as the Jordan River flows through an agricultural basin (^12^).

**Table S1.** Physicochemical characterization of the porewater in the hyporheic zone.

| **Season Parameter** |  | **Summer** | | | **Winter** | |
| --- | --- | --- | --- | --- | --- | --- |
|  | **Units** | **Aerobic** | **Suboxic** | **Aerobic** | | **Suboxic** |
| Depth | cm | 1 | 7 ± 6 | 11 ± 3.6 | | 14 ± 4 |
| Temperature | °C | 24 ± 1.3 | 30 ± 4.6 | 16 ± 1 | | 18 ± 1.5 |
| DO | mg L^-1^ | 8.1 ± 0.6 | 1.2 ± 0.8 | 9.6 ± 0.6 | | 2.6 ± 1.3 |
| Oxygen saturation | % | 83 ± 6.2 | 12 ± 7.6 | 93 ± 5 | | 26 ± 13 |
| NO_3_^-^ - N | µmol L^-1^ | 75 ± 63 | 6.6 ± 6.5 | 132 ± 22 | | 112 ± 34 |
| NH_4_^+^ - N | µmol L^-1^ | 2.8 ± 0.9 | 2.9 ± 0.9 | 0.8 ± 0.9 | | 1.1 ± 0.0 |
| PO_4_^3-^ - P | µmol L^-1^ | 98 ± 24 | 112 ± 53 | 61 ± 31 | | 66 ± 42 |
| N:P | mol:mol | 144 ± 118 | 8.7 ± 9.4 | 253 ± 144 | | 263 ± 106 |
| TOC | mg L^-1^ | 10 ± 3.7 | 24 ± 14 | 2.3 ± 1.5 | | 3.8 ± 3.4 |
| TON | mg L^-1^ | 2.1 ± 0.5 | 7.3 ± 0.6 | 2.4 ± 1.5 | | 1.1 ± 0.6 |
| C:N | mol:mol | 4.7 ± 1.0 | 3.5 ± 2.2 | 2.0 ± 0.9 | | 8.6 ± 6.7 |

**Table S2.** Physicochemical profile of the hyporheic zone in the Jordan River (32° 54' 21.4704'' N, 35° 37' 7.4712'' E) at Gesher Arik at antipodal seasons and across oxygen gradients. Stream profile of abiotic conditions and biofilm nutrient composition and stoichiometric ratios.

|  | **Season Parameter** | **Summer** | | **Winter** | |
| --- | --- | --- | --- | --- | --- |
|  |  | **Aerobic** | **Suboxic** | **Aerobic** | **Suboxic** |
| Stream characteristics | Depth, overlying water (cm) | 100 | — | 60 | — |
|  | Discharge flux (m^3^ sec^-1^) | 6.9 ± 0.4 | — | 12.9 ± 4.9 | — |
|  | EC (μS/cm^2^) | 410 ± 5 | 951 ± 602 | 514 ± 80 | 817 ± 321 |
|  | pH | 7.9 | 6.9 | 7.5 | 7.4 |
| Biofilm composition | DOC (mg C g^-1^) | 1.3 ± 0.8 | 0.8 ± 0.2 | 2.1 ± 0.9 | 2.3 ± 0.9 |
|  | TN (mg N g^-1^) | 1.4 ± 0.7 | 0.9 ± 0.2 | 0.6 ± 0.3 | 0.6 ± 0.3 |
|  | DN (mg N g^-1^) | 0.2 ± 0.0 | 0.2 ± 0.0 | 0.5 ± 0.2 | 0.6 ± 0.3 |
|  | C:N (mol:mol) | 4.1 ± 0.2 | 4.1 ± 0.3 | 5.1 ± 1.5 | 4.6 ± 1.0 |
|  | DC:DN (mol:mol) | 5.4 ± 2.9 | 4.5 ± 0.9 | 4.3 ± 1.0 | 4.3 ± 0.9 |

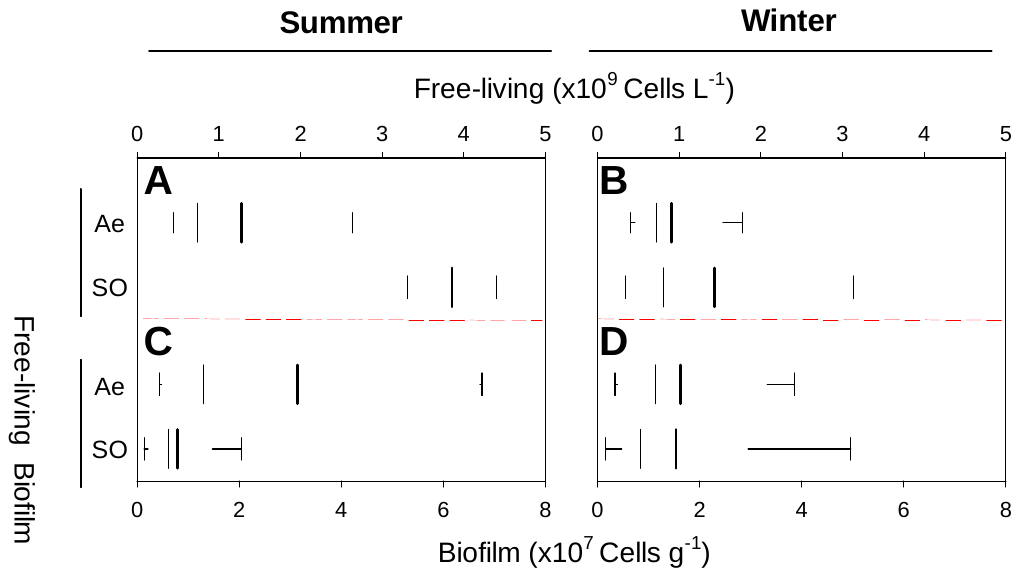

**Figure S2.** Bacterial abundance sampled from the hyporheic zone during summer (A, C) and winter (B, D). These values were measured at aerobic (Ae, cyan) and suboxic (SO, red) layers for free-living (pastel colors) and biofilm associated diazotrophs (diagonal hatching). Boxplots Ae FL represents n = 3 and 4 in summer and winter respectively, SO FL represents n = 2 and 3, Ae Bio represents n = 5 and 4, and SO Bio represents n = 5 and 5 in summer and winter, respectively. Box plots (A-D) portray interquartile range (25^th^ and 75^th^ percentiles) with whiskers extending to the 5^th^ and 95^th^ percentiles with a mean line (thick) and a median line (thin).

**Submission numbers of the study sequenced from the Jordan River**

Sequence Read Archive (SRA) submission: SUB16126134
Free-living and biofilm-associated riverbed diazotroph diversity, Apr 15 '26
BioProject: Processed
PRJNA1454053: Free-living and biofilm-associated riverbed diazotroph diversity

**Table S3.** Representative Order proportions and corresponding dominant genus within sample. Each sample was sequenced once, except for winter FL Ae and SO and Bio Ae that were sampled twice.

| Order | Genus | Summer Ae_FL | Summer SO_FL | Summer Ae_Bio | Summer SO_Bio | Winter Ae_FL | Winter SO_FL | Winter Ae_Bio | Winter SO_Bio |
| --- | --- | --- | --- | --- | --- | --- | --- | --- | --- |
| Bacillales | Paenibacillus | 19.77 | 0.0 | 0.46 | 0.0 | 10.04 | 2.44 | 0.61 | 1.12 |
| Bacteroidia* | * | 2.01 | 2.91 | 1.61 | 1.73 | 1.83 | 0.51 | 0.0 | 1.1 |
| Marinilabiliales | Draconibacterium | 0.0 | 0.0 | 2.21 | 1.1 | 0.61 | 0.51 | 0.0 | 0.77 |
| Cyanobacteria* | * | 0.0 | 0.0 | 1.61 | 0.33 | 0.0 | 0.0 | 0.0 | 1.82 |
| Cyanobacteria* | * | 0.0 | 0.0 | 0.0 | 0.0 | 0.0 | 0.0 | 0.57 | 0.27 |
| Cyanophyceae * | * | 1.17 | 3.62 | 0.46 | 1.73 | 0.53 | 5.28 | 0.0 | 0.04 |
| Myxococcia* | * | 5.53 | 2.36 | 0.55 | 0.4 | 8.63 | 5.33 | 0.0 | 1.56 |
| Nitrospirales | * | 1.94 | 2.83 | 0.64 | 1.1 | 0.0 | 0.0 | 0.08 | 1.26 |
| Nitrospirales | RPQD01000082,  OGW20163.1 | 0.0 | 0.0 | 1.01 | 1.1 | 0.53 | 0.0 | 2.11 | 0.9 |
| Nitrospirota* | * | 0.0 | 0.24 | 0.64 | 5.52 | 0.0 | 0.56 | 2.07 | 2.93 |
| SCSY01000135* | * | 0.0 | 0.0 | 0.18 | 0.7 | 0.0 | 0.93 | 0.53 | 0.06 |
| Alphaproteobacteria* | * | 0.84 | 1.81 | 0.0 | 0.23 | 0.44 | 0.27 | 0.0 | 0.0 |
| Hyphomicrobiales | Bradyrhizobium | 4.98 | 17.24 | 10.11 | 8.34 | 7.91 | 6.91 | 4.87 | 7.93 |
| Sphingomonadales | Novosphingobium | 0.0 | 0.87 | 0.0 | 0.0 | 1.03 | 0.77 | 4.06 | 1.83 |
| Betaproteobacteria incertae sedis | Candidatus Accumulibacter | 0.0 | 0.79 | 0.97 | 5.32 | 0.13 | 0.36 | 0.0 | 0.0 |
| Betaproteobacteria* | * | 0.59 | 0.0 | 1.56 | 0.76 | 0.21 | 0.63 | 1.01 | 3.33 |
| Burkholderiales | * | 5.12 | 10.31 | 13.69 | 5.15 | 5.21 | 6.35 | 15.82 | 3.73 |
| Rhodocyclales | Dechloromonas | 3.07 | 6.61 | 4.64 | 2.36 | 5.73 | 3.33 | 3.65 | 4.48 |
| Aeromonadales | Tolumonas | 34.96 | 1.73 | 3.22 | 0.0 | 15.13 | 3.94 | 0.0 | 0.0 |
| Enterobacterales | Klebsiella | 0.37 | 1.65 | 0.46 | 0.57 | 0.11 | 0.46 | 0.0 | 0.95 |
| Gammaproteobacteria incertae sedis | Candidatus Competibacteraceae* | 0.0 | 0.0 | 0.6 | 0.0 | 0.52 | 0.42 | 0.69 | 0.35 |
| Gammaproteobacteria* | * | 3.7 | 3.23 | 6.85 | 3.46 | 2.72 | 3.24 | 11.27 | 12.8 |
| Vibrionales | Vibrio | 0.0 | 0.0 | 1.65 | 3.76 | 0.55 | 0.0 | 4.87 | 2.69 |
| Pseudomonadota* | * | 3.0 | 4.33 | 4.92 | 8.94 | 2.89 | 2.78 | 12.49 | 8.66 |
| Hyphomicrobiales | * | 1.9 | 2.68 | 1.33 | 0.53 | 1.32 | 1.6 | 0.0 | 1.04 |
| Hyphomicrobiales | Methylocystis | 0.0 | 0.0 | 2.99 | 0.63 | 0.0 | 0.63 | 1.05 | 0.37 |
| Candidatus Dadabacteria* | * | 0.0 | 2.91 | 2.21 | 2.86 | 2.99 | 13.75 | 2.55 | 0.51 |
| Desulfobacteria* | * | 0.37 | 0.0 | 0.37 | 0.5 | 1.12 | 0.08 | 0.0 | 0.22 |
| Desulfobulbaceae | * | 1.72 | 1.34 | 1.1 | 2.49 | 1.66 | 0.46 | 0.0 | 0.0 |
| Desulfobulbales | Desulfobulbus | 1.94 | 4.57 | 3.77 | 2.06 | 0.55 | 0.62 | 1.99 | 0.0 |
| Desulfovibrionales | Oceanidesulfovibrio | 0.0 | 6.38 | 6.3 | 1.16 | 0.31 | 5.62 | 0.77 | 1.57 |
| Desulfuromonadales | Geobacter | 2.75 | 19.21 | 10.06 | 8.48 | 8.9 | 12.42 | 21.82 | 17.59 |
| Desulfuromonadia* | * | 0.0 | 0.0 | 4.37 | 12.9 | 0.0 | 0.76 | 1.91 | 0.0 |
| Thermodesulfobacteriota* | * | 3.66 | 1.26 | 5.79 | 11.5 | 7.01 | 16.72 | 4.46 | 13.17 |
| Verrucomicrobiota* | * | 0.62 | 1.1 | 3.68 | 4.29 | 11.39 | 2.33 | 0.77 | 6.96 |

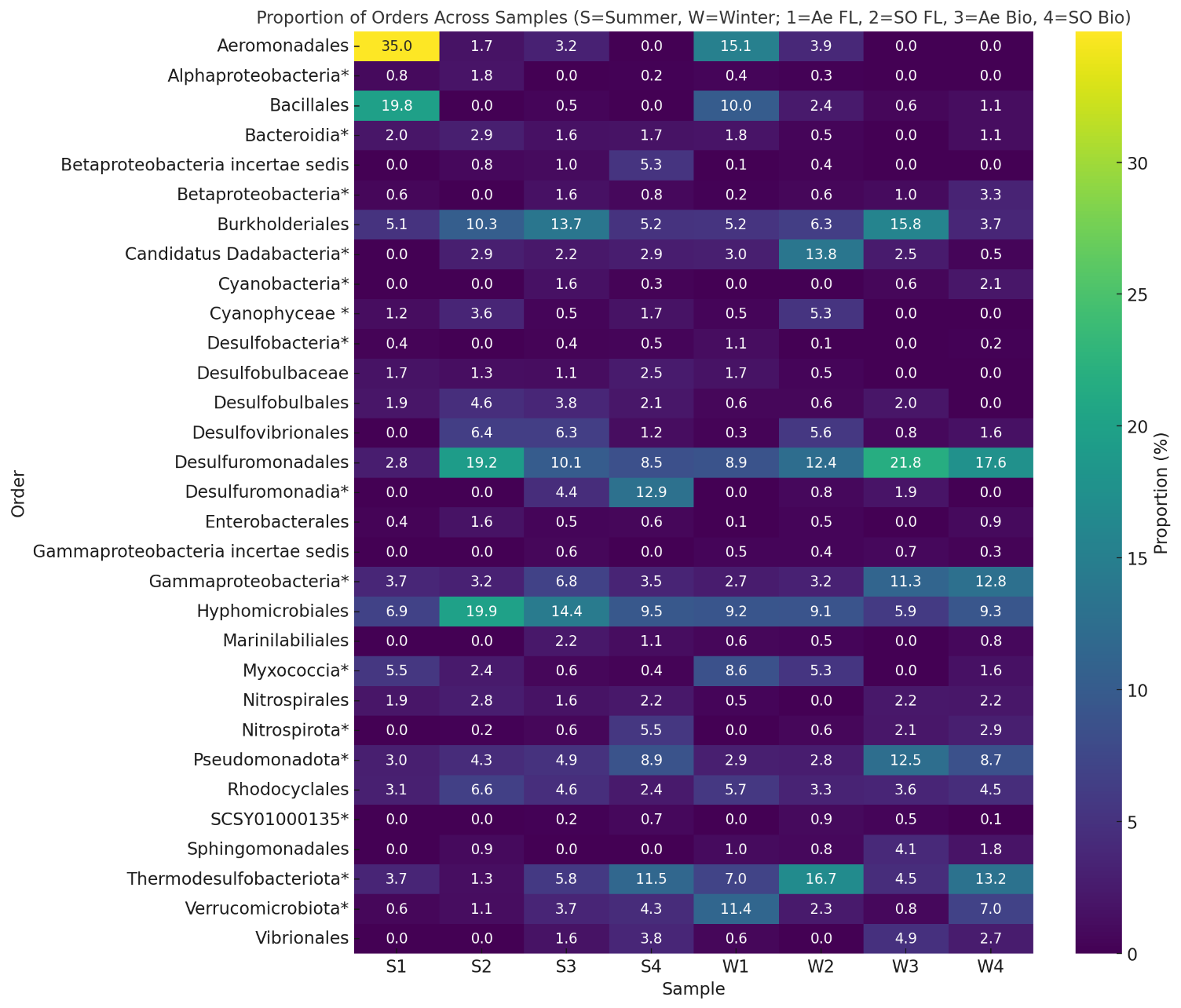

**Figure S3.** Overall proportions of orders of sequenced *nifH* within samples from summer and winter, free-living and biofilm-associated, and aerobic and suboxic zones.

**Table S4.** Parameters for biofilm-associated diazotroph normalization to volume calculation.

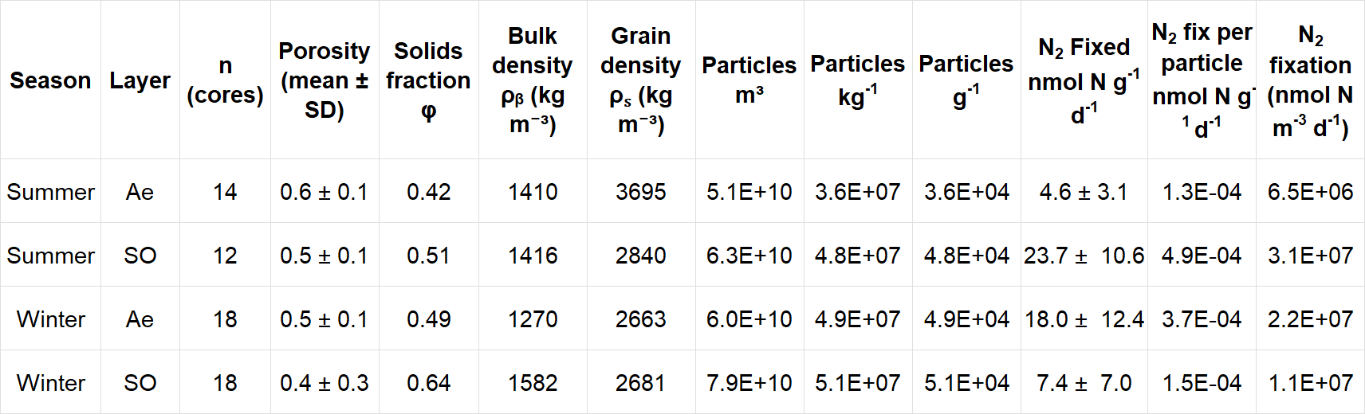

**Table S5**. N_2_ fixed volumetrically in nmol N m^-3^ d^-1^ compared to global reports from rivers and lakes in the water column and benthos and local rates in the water column of Gesher Arik. Water column N_2_ fixation rates are sourced from Geisler et al. (^3^) and global reports are from Fulweiler et al. (^13^). N_2_ fixation rates from the sampling site are the averaged and summed rates from the water column (free-living and aggregate-associated) and the hyporheic zone (free-living and biofilm-associated) in summer and winter. The global reports are the average rates that were measured volumetrically.

| Season | Layer | Lifestyle | | Overlying Water | | Lake | | River | |
| --- | --- | --- | --- | --- | --- | --- | --- | --- | --- |
|  |  | Bio  (n=23) | FL  (n=24) | Agg  (n=12) | FL  (n=8) | WC (n=462) | Benthos (n=11) | WC (n=36) | Benthos (n=1) |
| Summer | Ae | 6.49 × 10⁶ (4.4× 10^6^) | 0 | 246.6 (79.6) | 40.1 (6.4) | 6.10 × 10⁴ | 7.51 × 10⁶ | 1.29 × 10⁵ | 0 |
|  | SO | 3.12 × 10⁷ (1.4 × 10^7^) | 7.70 × 10³ (9.3× 10^3^) | – | – | – | – | – | – |
| Winter | Ae | 2.20 × 10⁷ (1.5× 10^7^) | 3.31 × 10³ (1.9× 10^3^) | 104.2 (42.2) | 83.3 (70.4) | – | – | – | – |
|  | SO | 1.15 × 10⁷ (1× 10^7^) | 3.07 × 10³ (3.3× 10^3^) | – | – | – | – | – | – |
